# Synthetic collinearization of a chromosome segment enables genetic dissection of incompatibility between distant genera

**DOI:** 10.1101/2025.09.29.679105

**Authors:** Shawn H. Yang, Alessandro L. V. Coradini, Cara B. Hull, Daniel T. Lusk, Ian M. Ehrenreich

## Abstract

Reproductive barriers and genomic structural differences hinder genetic analysis between highly divergent organisms. Here, we examined whether these challenges can be overcome by collinearization, the synthesis of chromosomes that retain the native gene content and organization of a host organism while incorporating the DNA sequences of another organism. We applied collinearization to *Kluyveromyces marxianus*, a yeast species that is both distantly related to and has a distinct genome structure from the model yeast *Saccharomyces cerevisiae*. We generated a ∼35-kb *K. marxianus* DNA segment that was collinear with one-sixth of a *S. cerevisiae* chromosome and contained 17 protein-coding genes. Although this synthetic chromosome segment successfully substituted for its native counterpart in an *S. cerevisiae* cell, it imposed a significant growth cost due to the incompatibility of two *K. marxianus* proteins with the host proteome. We predicted the disrupted protein-protein interactions using AlphaFold and alleviated their cost by supplementing orthologous protein partners from *K. marxianus*. Furthermore, we completely eliminated the growth cost by replacing the two incompatible *K. marxianus* genes with their *S. cerevisiae* orthologs. These findings demonstrate that collinearization can facilitate the creation of chimeric genomes to dissect the genetic basis of incompatibility and traits between organisms that cannot naturally hybridize. They also suggest that hundreds of genetic incompatibilities exist between *S. cerevisiae* and *K. marxianus*, reflecting disrupted protein-protein interactions that may be predictable *in silico*.

## Introduction

The remarkable diversity of organisms in nature provides an invaluable resource for understanding the molecular mechanisms underlying the evolution of species and traits ^1–3^. However, the most effective genetic methods for investigating the basis of this diversity, such as linkage mapping and genome-wide association studies, are largely restricted to use within species or between closely related species ^4^. As the evolutionary distance between organisms increases, the likelihood of reproductive barriers and genome structural differences grows substantially ^5–8^, making the application of these conventional genetic methods increasingly challenging.

The synthesis of chromosomes and genomes, known as synthetic genomics, can help overcome these obstacles. Over the past ∼15 years, advances in synthetic genomics have enabled the *de novo* synthesis of large DNA molecules, ranging from tens of kilobases to multiple megabases ^9,10^. Synthetic genomics makes possible the complete reprogramming of the sequence and structure of these molecules, opening new avenues for investigating fundamental questions in biology. For example, recent studies have used synthetic genomics to examine the minimal set of genes required to produce a bacterial cell ^11^, the plasticity of the genetic code ^12^, and the principles of mammalian *Hox* cluster regulation ^13^.

In the context of evolutionary biology, synthetic genomics may facilitate the genetic dissection of traits and isolating mechanisms between divergent organisms that cannot interbreed or produce viable offspring ^9^. However, the differences in genome content and structure that abound across species are a barrier to such work. In this paper, we explored whether synthetic genomics can be used to artificially establish synteny between, or ‘collinearize’ ^14^, the genomes of highly divergent organisms, thereby enabling genetic analyses that would not otherwise be possible. For example, collinearization can improve understanding of how incompatibilities accumulate between organisms over vast evolutionary time, which at present is difficult to address experimentally on a genomic scale ^5–8^.

To explore the potential of collinearization, we focused on two distantly related budding yeast species that cannot naturally hybridize: *Saccharomyces cerevisiae* and *Kluyveromyces marxianus*. Although both species have genomes of ∼12 Mb containing ∼5,000 genes, their genome structures are entirely distinct–e.g., *S. cerevisiae* has 16 chromosomes, whereas *K. marxianus* has 8. In our collinearization study, we used *S. cerevisiae* as our genomic template and host cell, and *K. marxianus* as our source of DNA sequences. We conducted this work in *S. cerevisiae* because it is a well-established model organism, including for synthetic genomics ^15^ and research on the genetic basis of isolating mechanisms between yeast strains and species ^16,17,14,18–25^. We chose *K. marxianus* because it is widely used in biotechnology for its fast growth rate and exceptional thermotolerance and pH tolerance ^26^.

Here, we designed a complete chromosome *in silico* that preserved the native gene content and organization of *S. cerevisiae* while replacing all its DNA sequences with those from *K. marxianus*. As proof of concept, we then constructed the first one-sixth of this synthetic *K. marxianus* chromosome, and analyzed its transcription and phenotypic effects in an *S. cerevisiae* cell. We found that this segment was initially incompatible with the *S. cerevisiae* cellular environment and identified the causative genes and disrupted protein–protein interactions. Addressing these incompatibilities allowed us to produce a fully compatible synthetic molecule.

## Results

### Design of synKmI, a K. marxianus chromosome collinear to S. cerevisiae ChrI

*S. cerevisiae and K. marxianus* exhibit significant divergence in nucleotide sequence, protein sequence, and genome structure. On average, coding regions show 44% nucleotide divergence and 49% amino acid divergence, with 26% of aligned nucleotide positions containing gaps (Supplementary Fig. 1). Noncoding regions show higher nucleotide divergence, with promoter regions displaying around 61% nucleotide divergence and 50% of aligned nucleotide positions containing gaps (Supplementary Fig. 2). The Kluyveromyces and Saccharomyces lineages diverged before a whole-genome duplication (WGD) event that occurred approximately 114 million years ago (Fig. 1a) ^27,28^. The WGD took place in the lineage that gave rise to Saccharomyces, followed by a reduction in genome content that resulted in similar gene content but minimal genomic correspondence between *S. cerevisiae* and *K. marxianus* ^29^. For instance, orthologous regions of *S. cerevisiae* Chromosome I (ChrI) are dispersed across synteny blocks on seven of the eight *K. marxianus* chromosomes (Fig. 1b).

**Figure 1.**
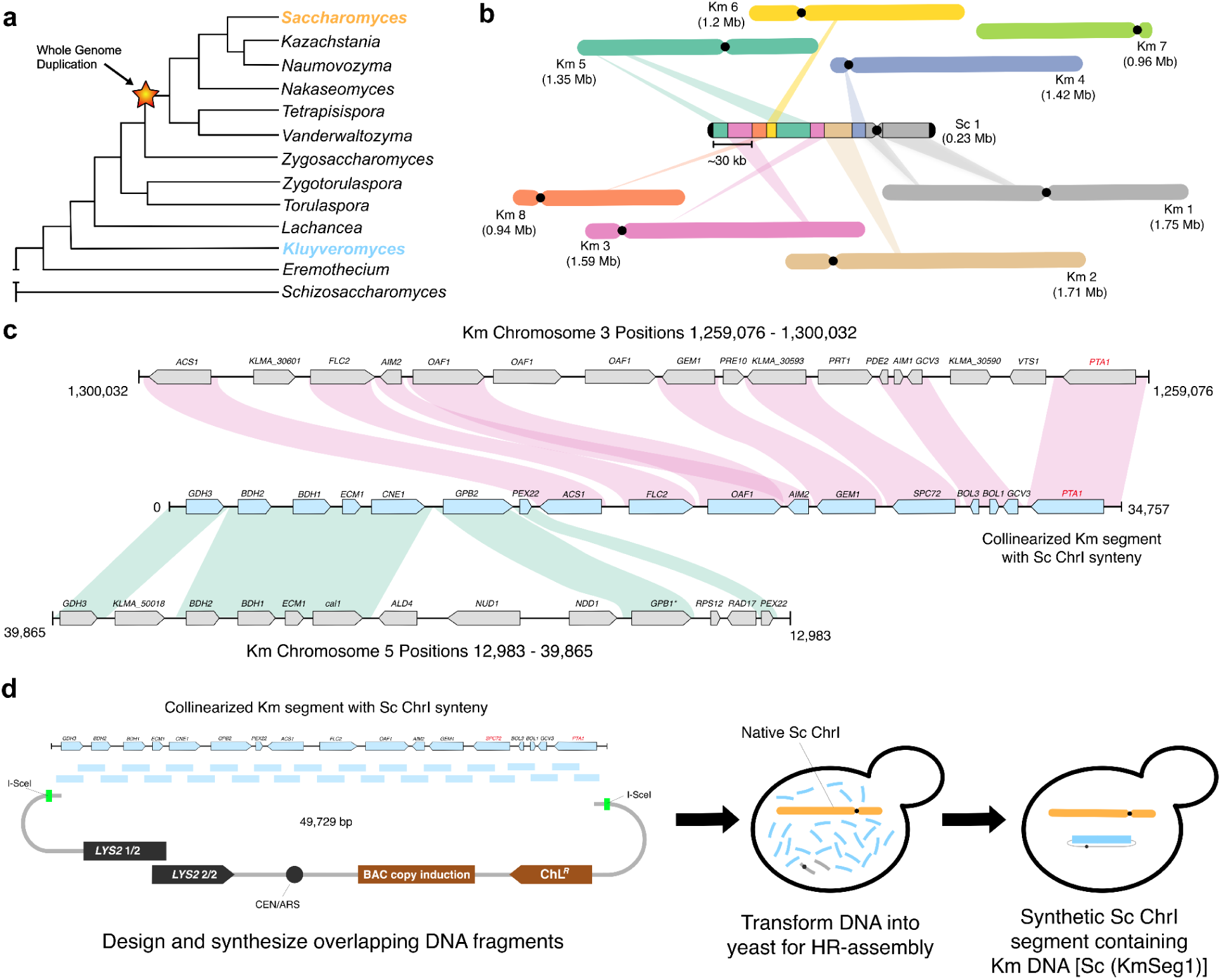
Design and synthesis of a collinearized chromosome segment across genera. **a.** The *Kluyveromyces* genus predates a whole genome duplication that occurred prior to the formation of the *Saccharomyces* genus. The phylogeny of the yeast genera is based on ^49,50^. **b.** Genome structure varies significantly between *Kluyveromyces marxianus* (Km) and *Saccharomyces cerevisiae* (Sc). Synteny blocks between *S. cerevisiae* Chromosome I (ChrI) and the *K. marxianus* genome were identified and mapped across seven out of the eight *K. marxianus* chromosomes. **c.** Orthologous regions from *K. marxianus* were used to design a *K. marxianus* segment that was collinear to *S. cerevisiae* ChrI. Here, we show the first of six segments of this collinear chromosome, which we refer to as KmSeg1. **d.** Overlapping DNA fragments were designed and synthesized for in vivo assembly via homologous recombination in yeast, resulting in a collinear *K. marxianus* KmSeg1 inside an *S. cerevisiae* cell.

We designed *in silico* a *K. marxianus* chromosome, which we call ‘synKmI,’ that is collinear with *S. cerevisiae* Chromosome I (ChrI), using nucleotide sequences—both coding and noncoding—exclusively from *K. marxianus* (Fig. 1c; Supplementary Figs. 3 and 4). We started this design from the beginning of the left subtelomeric region of *S. cerevisiae* ChrI and added *K. marxianus* sequences to it until we reached the end of the right subtelomeric region of ChrI ^30^. The only exceptions were the centromere, where we used the *S. cerevisiae* ChrI centromere sequence (*CEN1*) to ensure proper chromosome maintenance, and any mobile genetic elements, which were excluded from the design. These choices were made to ensure that our insights would not be influenced by abnormal chromosome segregation ^31^, as recent work has shown that *K. marxianus* centromeres are not effective in *S. cerevisiae* ^32^.

When possible, sets of consecutive genes from the *K. marxianus* genome were incorporated without modification if their gene content and orientation exactly matched the *S. cerevisiae* genome (Fig. 1c). In such cases, the full intergenic regions between terminal genes and middle genes were included. For terminal genes at the boundaries of these sets, the promoter was defined as the shorter of either 500 bp upstream of the start codon or the entire intergenic region ^33^. In cases where the terminator of a terminal gene was at the end of a set, we defined the terminator as 100 bp downstream of the stop codon or the full intergenic region, whichever was shorter. In *K. marxianus*, 3′ UTRs have not been comprehensively defined, so determining how to handle regions downstream of genes from this species was necessarily subjective. The most common 3′ UTR length in *S. cerevisiae* is under 100 nucleotides ^34^ and numerous synthetic terminators have been designed successfully with lengths of 35 to 70 nucleotides ^35^. Based on this, we selected 100 nucleotides as the terminator for cases where a gene was the terminal gene in a set.

Many *K. marxianus* genes were not located in sets of consecutive genes. In these cases, the *K. marxianus* gene, along with its promoter and terminator sequences, was appended to the design of synKmI using the same approach described above. Each *K. marxianus* gene, along with its promoter and terminator, was oriented as in *S. cerevisiae*. In addition, since *K. marxianus* is a pre-WGD species, some genes present as paralogs in *S. cerevisiae* exist as a single copy in *K. marxianus*; in such cases, we included the single *K. marxianus* gene. In rare instances of tandem gene duplications in *K. marxianus* that are absent in *S. cerevisiae*, we selected the copy with the highest homology to the *S. cerevisiae* ortholog, determined using Protein BLAST ^36^. By systematically applying these rules, we ensured that synKmI was collinear with *S. cerevisiae* ChrI and contained the maximal amount of *K. marxianus* sequences.

### Synthesis and expression of synKmI segment 1 (KmSeg1)

Due to concerns about potential gene or protein misexpression or genetic incompatibilities rendering synKmI nonviable, we synthesized only 1/6th of it: a ∼35-kb segment containing 17 protein-coding genes, including a single essential gene, *PTA1* (Fig. 1c). Based on the *in silico* design, we ordered this segment, referred to as synKmI segment 1 (abbreviated hereafter ‘KmSeg1’), as 25 partially overlapping synthetic DNA fragments, each up to 1.8 kb in size (Fig. 1d). To facilitate assembly of all the fragments by homologous recombination in yeast, each fragment was synthesized with at least 200 bp of homology to its adjacent fragment(s) or the pASC1 BAC/YAC vector ^37^. In this assembly, we provided pASC1 as two partially overlapping fragments, each containing a portion of the *LYS2* marker, resulting in a total of 27 DNA molecules.

The fragments were assembled with each other and pASC1 in a single transformation of the *S. cerevisiae* BY4742 strain. The transformation yielded ∼300 colonies, of which 8 were screened using PCR checks at a subset of assembly junctions (Supplementary Fig. 5). All examined colonies exhibited the correct PCR amplicons, indicating highly efficient and accurate assembly. We selected one colony for Oxford Nanopore long-read sequencing, which confirmed that the sequence and structure of KmSeg1 matched its *in silico* design exactly (Supplementary Fig. 6).

We assessed whether *K. marxianus* genes on KmSeg1 are expressed in *S. cerevisiae*. Given the substantial divergence between these species, *K. marxianus* promoters might not drive transcription in *S. cerevisiae*. However, we found that all *K. marxianus* genes on segment 1 were transcribed in an *S. cerevisiae* cell (Fig. 2a), with transcription occurring largely within the expected gene boundaries. Using RNA-seq data from the *S. cerevisiae* strains containing both the native ChrI and the synthetic segment 1, we directly compared the expression of *K. marxianus* genes to their respective *S. cerevisiae* orthologs. Five *K. marxianus* genes—*ACS1*, *AIM2, BDH1, BDH2,* and *FLC2*—were expressed at significantly higher or lower levels than their *S. cerevisiae* counterparts (paired t-test, *p* < 0.05, Benjamini-Hochberg FDR < 0.1; Fig. 2b), with differences ranging from ∼8-fold to ∼512-fold. These differences in gene expression between *K. marxianus* and *S. cerevisiae* alleles of the same genes likely reflect variation in *cis*-regulatory elements between the species.

**Figure 2.**
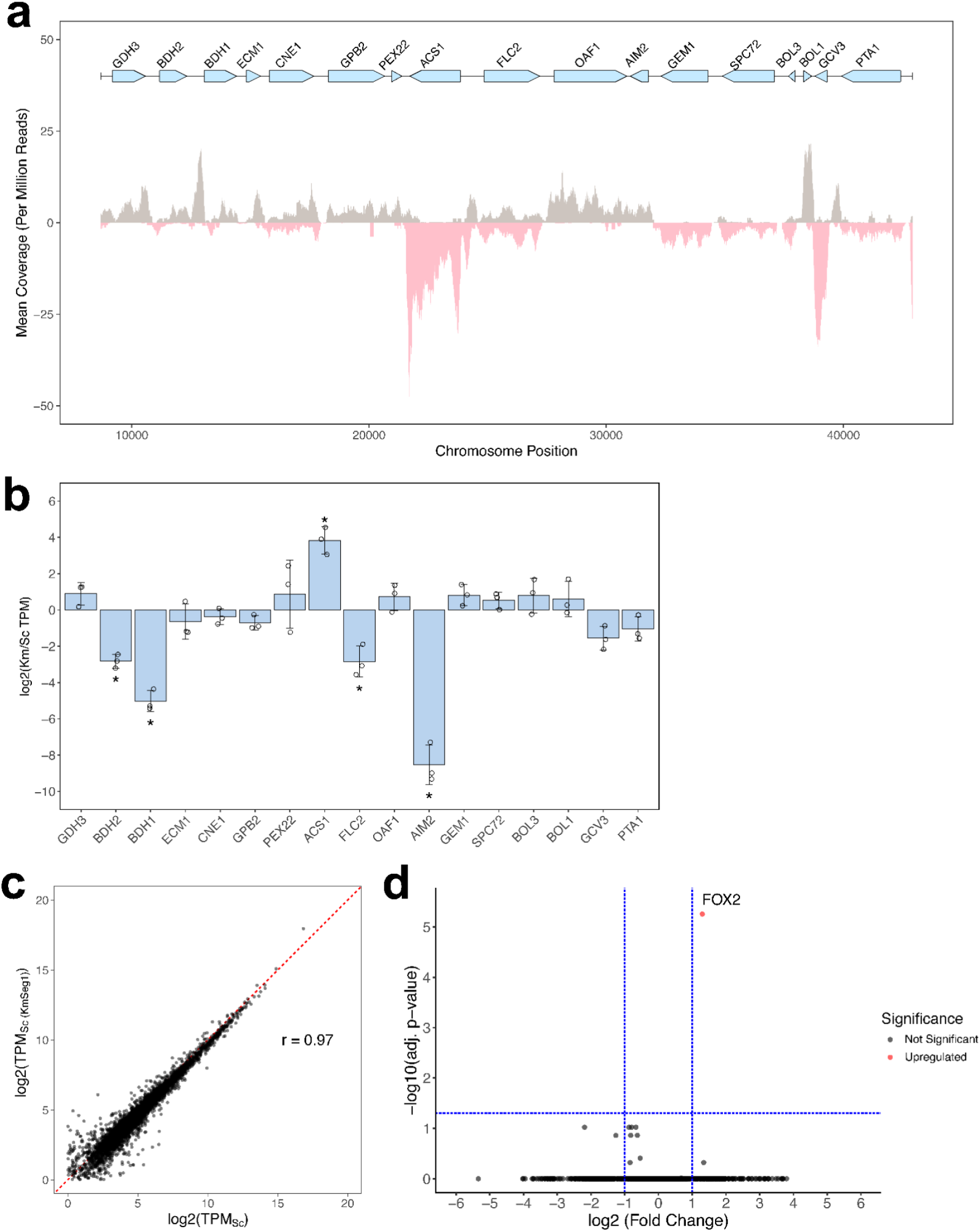
Transcriptomic analysis of KmSeg1 in an *S. cerevisiae* cell. **a.** Strand-specific RNA-seq coverage plot of KmSeg1 in an *S. cerevisiae* cell. Coverage of KmSeg1 is displayed as the mean coverage per million reads across three biological replicates (n = 3). Plus and minus strand coverage is represented by gray and pink, respectively. **b.** Comparison of ortholog expression levels within the same *S. cerevisiae* cell. The y-axis represents the log_2_ ratio of transcripts per million (TPM) values (log_2_(TPM_K. marxianus_/TPM_S. cerevisiae_)). Asterisks denote significant differences in expression between *S. cerevisiae* and *K. marxianus* alleles in the same cell (paired t-test, *p* < 0.05, Benjamini-Hochberg FDR < 0.1). **c.** Genome-wide transcription remains largely unaffected in the presence of *K. marxianus* DNA. Scatterplot showing strongly correlated genome-wide gene expression levels of the *S. cerevisiae* progenitor strain (x-axis) versus the *S. cerevisiae* strain harboring KmSeg1 (y-axis; Pearson correlation, r = 0.97, *p* ≅ 0). **d.** Volcano plot of genome-wide gene expression in the *S. cerevisiae* control strain and the *S. cerevisiae* strain with KmSeg1. The only gene with significant upregulation is *FOX2* and no genes were seen with significant downregulation (Wald test, Benjamini-Hochberg FDR < 0.05).

We also compared global gene expression between *S. cerevisiae* strains with and without KmSeg1. For the rest of the transcriptome beyond KmSeg1, gene expression showed a strong correlation (Pearson correlation, r = 0.97, *p* ≅ 0; Fig. 2c). The only gene with significant upregulation was *FOX2* and no genes were seen with significant downregulation (Wald test, Benjamini-Hochberg FDR < 0.05; Fig. 2d). This result shows that genome-wide transcription in a wild type *S. cerevisiae* cell remains largely unaffected in the presence of KmSeg1.

### KmSeg1 substitutes for its S. cerevisiae counterpart

After confirming that KmSeg1 is actively transcribed in *S. cerevisiae*, we next tested whether it could substitute for its native counterpart and support cell viability. Using CRISPR-Cas9, we deleted the native segment on *S. cerevisiae* ChrI and replaced it with a *URA3* marker (Fig. 3a). Transformants were successfully recovered by selecting for uracil prototrophy, confirming the absence of the native segment (Supplementary Fig. 7). Because *PTA1* is essential in *S. cerevisiae*, the recovery of this deletion strain indicates that *K. marxianus PTA1* is functional and capable of complementing its native ortholog. This result suggests that other *K. marxianus* genes in KmSeg1 can function in an *S. cerevisiae* cellular environment.

**Figure 3.**
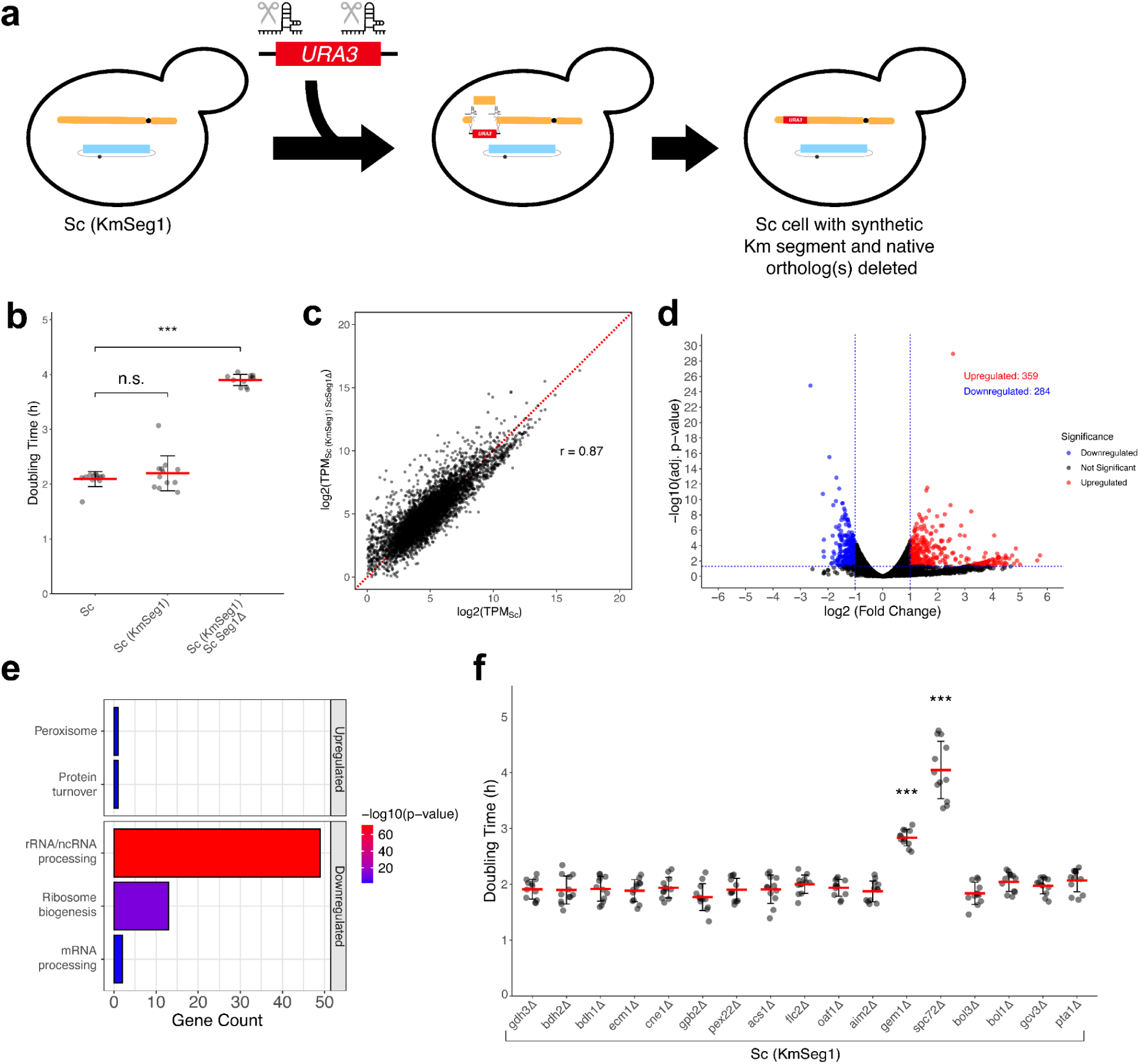
Analysis of KmSeg1 in an *S. cerevisiae* cell after deletion of its native counterpart. **a.** Schematic overview of the process to delete the native *S. cerevisiae* counterpart of KmSeg1. In the *S. cerevisiae* cell harboring KmSeg1, gRNAs targeting both ends of the targeted region, along with a selectable *URA3* repair template, are transformed into a donor cell constitutively expressing Cas9. Transformants were then screened for correct insertion of the deletion cassette and euploidy using PCR checks. **b.** Substitution of KmSeg1 causes *S. cerevisiae* to show a growth defect. Red bars represent the mean doubling time, and error bars indicate the standard deviation calculated from four biological replicates and three technical replicates each (n = 12). Statistical significance was assessed using a two-tailed t-test (*** denotes *p* < 0.001 and n.s. denotes *p* > 0.05). **c.** Genome-wide transcription is disrupted by substitution of KmSeg1 for its native *S. cerevisiae* counterpart. Scatterplot of genome-wide gene expression levels in the *S. cerevisiae* control strain (x-axis) versus the *S. cerevisiae* strain with the KmSeg1 substitution (y-axis; Pearson correlation, r = 0.87). **d.** Volcano plot showing differentially expressed genes between the *S. cerevisiae* control strain and the *S. cerevisiae* strain with the KmSeg1 substitution. Upregulated genes are shown in red and downregulated genes are shown in blue (Wald test, Benjamini-Hochberg FDR < 0.05). **e.** Functional analysis of differentially expressed genes. Upregulated (top) and downregulated (bottom) genes identified from differential expression analysis were used as input for spatial analysis of functional enrichment via TheCellMap^38^. Enrichments with *p* ≤ 0.05 are displayed using barplots. **f.** Two *K. marxianus* genes, *GEM1* and *SPC72*, are responsible for the growth defect caused by the KmSeg1 substitution. Individual deletions of native *S. cerevisiae* orthologs were tested in the presence of KmSeg1, identifying the two incompatible *K. marxianus* genes. One-way ANOVA (*p* = 3.15 × 10-^70^) followed by Tukey’s Honest Significant Difference test (95% confidence level) was performed (*** denotes *p* < 0.001 and no asterisk denotes *p* > 0.05).

An *S. cerevisiae* strain containing both KmSeg1 and its native counterpart showed no growth defect (difference between *S. cerevisiae* with *K. marxianus* segment vs. *S. cerevisiae* control = 6.37 ± 20.79 minutes [mean ± 1 s.d. hereafter], t-test, *p* > 0.05). However, substitution of KmSeg1 for its native counterpart results in a significant growth defect, with a ∼2-fold increase in doubling time (difference between *K. marxianus* segment substitution and *S. cerevisiae* control = 108.44 ± 10.18 minutes, t-test, *p* = 3.78 x 10^-20^; Fig. 3b). These results indicate that the KmSeg1 has a recessive deleterious effect in *S. cerevisiae*, which is masked when the native copy of this segment is present but becomes apparent when the native copy is deleted.

Following deletion of the native segment, significant genome-wide expression differences emerged relative to the *S. cerevisiae* control that were not observed previously (Fig. 3c,d). When analyzing the differential expression of genes, 359 genes were upregulated, and 284 were downregulated (Wald test, Benjamini-Hochberg FDR < 0.05). To determine if the dysregulated genes were enriched for particular cellular processes, we utilized spatial analysis of functional enrichment via TheCellMap, leveraging the genetic interaction network of yeast (Fig. 3e, Supplementary Fig. 8) ^38^. Upregulated genes were enriched for peroxisome (*p* = 0.024) and protein turnover (*p* = 0.024), while downregulated genes were enriched for rRNA/ncRNA processing (*p* = 6.52 x 10^-72^), ribosome biogenesis (*p* = 1.53 x 10^-12^), and mRNA processing (*p* = 0.012). Although these functional enrichments suggest molecular mechanisms that may contribute to the growth defect observed upon substituting KmSeg1 for its native counterpart, they could also reflect a stress response resulting from the growth defect. The current data do not allow us to distinguish between these possibilities.

### Two genes on KmSeg1 are incompatible with an S. cerevisiae cell

To identify the specific gene(s) on KmSeg1 that were responsible for the growth defect, we generated single-gene deletions for all 17 *S. cerevisiae* orthologs within the strain containing KmSeg1, using the same approach employed to delete the entire region from ChrI (Fig. 3f). Among the deletions, two produced a recessive growth defect, while the other 15 had no effect (one-way ANOVA, *p* = 3.15 x 10^-70^, followed by Tukey’s Honest Significant Difference test with 95% family-wise confidence level).

One of the genes causing a growth defect was *GEM1*, which is known to be non-essential and encodes an outer mitochondrial membrane GTPase ^39^. Gem1 is a component of the ER-mitochondria encounter structure (ERMES) complex, which is involved in tethering the mitochondria and endoplasmic reticulum (ER) ^40^. We found that deletion of *GEM1* in wild type *S. cerevisiae* had no phenotypic effect (t-test, *p* > 0.05; Supplementary Fig. 9). The other gene causing a growth defect was *SPC72*, which encodes a γ-tubulin small complex (γ-TuSC) receptor and spindle pole body (SPB) component ^41^. We were unable to recover an *SPC72* deletion in wild type *S. cerevisiae*, suggesting that although *SPC72* is not annotated as essential, it is at least quasi-essential under our conditions ^39^.

The growth defects caused by *K. marxianus GEM1* and *SPC72* may result from coding variants, noncoding variants, or both. To identify whether the promoter or coding regions of *K. marxianus GEM1* and *SPC72* cause incompatibility with an *S. cerevisiae* cellular environment, we engineered chimeric versions of each gene by shuffling their promoter and coding regions between the genera (Supplementary Fig. 10). In these experiments, we defined promoters as 500 bp upstream of a start codon. For both genes, we generated all four possible promoter-coding region combinations as yeast centromeric plasmids (*S. cerevisiae*-*S. cerevisiae*, *K. marxianus*-*S. cerevisiae*, *S. cerevisiae*-*K. marxianus*, *K. marxianus*-*K. marxianus*). We then tested the phenotypic effect of each construct by transforming it into *S. cerevisiae* BY4742, deleting the native ortholog of *GEM1* or *SPC72*, and measuring the growth of the resulting strain.

For both *GEM1* and *SPC72*, the growth defect was due to the coding region, indicating that these proteins are incompatible with a *S. cerevisiae* cellular environment (t-test of constructs with *K. marxianus* coding region vs. *S. cerevisiae* control, *p* ≤ 3.20 x 10^-8^; Fig. 4a, b). However, our results indicate that the detrimental effects of *K. marxianus GEM1* are exacerbated when an *S. cerevisiae* promoter is used (t-test of construct with *S. cerevisiae* promoter and *K. marxianus* coding region vs. construct with *K. marxianus* promoter and *K. marxianus* coding region, *p* = 8.48 x 10^-7^). This finding suggests that the *GEM1* and *SPC72* promoters harbor *cis*-regulatory differences not detected by RNA-seq, and that these differences modestly influence the severity of the growth defect linked to each gene’s coding region.

**Figure 4.**
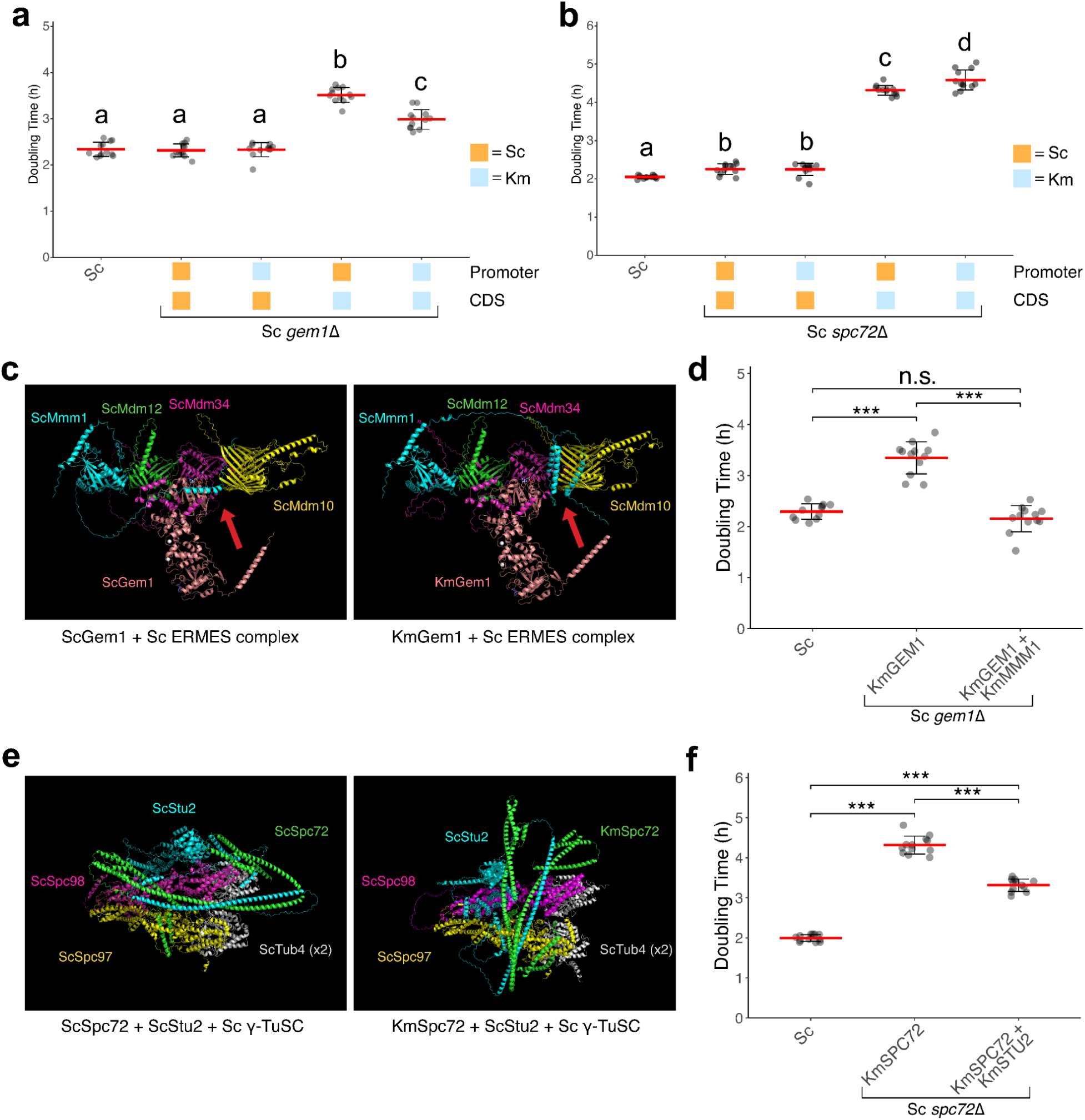
The proteins encoded by *K. marxianus GEM1* and *SPC72* are incompatible with *S. cerevisiae*. Dividing each gene into its promoter and coding region, we engineered the four possible chimeric versions of *GEM1* (**a**) and *SPC72* (**b**), identifying *K. marxianus* coding regions as the main source of incompatibility. Chimeric *GEM1* and *SPC72* constructs were tested in *S. cerevisia*e *gem1*Δ and *spc72*Δ backgrounds, respectively. Red bars indicate the mean doubling time from four biological replicates with three technical replicates each (n = 12). Error bars represent the standard deviation from the mean. Columns sharing the same letters are not significantly different (ɑ = 0.05, One-way ANOVA and Tukey’s HSD). **c.** Protein-protein interaction analysis of Gem1 and ERMES complex reveal a structural deformity in Mmm1 introduced by *K. marxianus* Gem1. Protein complex structures were predicted by AlphaFold3. Red arrows indicate the affected Mmm1 transmembrane domain. **d.** Supplementation of *K. marxianus* Mmm1 fully rescues the incompatibility observed with *K. marxianus* Gem1 in the *S. cerevisiae gem1*Δ background. Statistical significance was determined using using a two-tailed t-test (*** denotes *p* < 0.001 and n.s. denotes *p* > 0.05). **e.** Protein-protein interaction analysis of Spc72 and γ-tubulin small complex (γ-TuSC) reveals a structural deformity in Stu2 introduced by *K. marxianus* Spc72. **f.** Supplementation of *K. marxianus* Stu2 partially rescues the incompatibility observed with *K. marxianus* Spc72 in the *S. cerevisiae spc*72Δ background. Statistical significance was determined using using a two-tailed t-test (*** denotes *p* < 0.001 and n.s. denotes *p* > 0.05).

### GEM1 and SPC72 incompatibilities result from protein-protein interactions

Gem1 and Spc72 interact with multiple partner proteins, suggesting that the observed incompatibilities arise from disrupted protein-protein interactions. Since their divergence, *S. cerevisiae* and *K. marxianus* have likely accumulated amino acid differences that impair the physical interactions between *K. marxianus* Gem1 and Spc72 and their partner proteins in *S. cerevisiae.* To investigate this possibility, we employed AlphaFold3 to predict how substituting *K. marxianus* Gem1 and Spc72 for their *S. cerevisiae* counterparts might impact how these proteins complex with their *S. cerevisiae* partners ^42^. We then looked for structural abnormalities in the complexes that might indicate which partners contribute to the incompatibilities.

Gem1 is a component of the ERMES complex, which in yeast includes Mmm1, Mdm10, Mdm12, and Mdm34 ^43^. We modeled the *S. cerevisiae* alleles of these five proteins as a complex and then substituted *S. cerevisiae* Gem1 with *K. marxianus* Gem1 *in silico*. The predicted structure of the ERMES complex before and after the substitution indicated that *K. marxianus* Gem1 disrupts the structure of Mmm1 but not the other partners. Specifically, Mmm1 exhibited a structural deformity in a key transmembrane domain that anchors the ERMES complex to the ER (Fig. 4c). AlphaFold3 further predicted that this deformity could be rescued by replacing *S. cerevisiae* Mmm1 with *K. marxianus* Mmm1 when *K. marxianus* Gem1 is present (Supplementary Fig. 11). To test this, we supplemented *S. cerevisiae* BY4242 carrying the *S. cerevisiae*-*K. marxianus* (promoter-coding region) *GEM1* construct as centromeric plasmid and lacking the native *GEM1* with *K. marxianus MMM1* on an additional centromeric plasmid. We found that this fully rescued the incompatibility associated with *K. marxianus GEM1* (difference between supplemented strain vs. *S. cerevisiae* control = 8.00 ± 17.55 minutes, t-test *p* > 0.05; difference between supplemented strain vs. strain with *K. marxianus GEM1* growth defect = 70.95 ± 24.20 minutes, t-test *p* = 1.41 x 10^-9^; Fig. 4d). These results suggest that between *S. cerevisiae* and *K. marxianus*, *GEM1* and *MMM1* are involved in a Dobzhansky-Muller incompatibility ^44,45^.

Spc72 is a spindle pole body (SPB) component essential for proper spindle orientation during mitosis. Its N-terminus contains distinct binding sites for Stu2 and γ-TuSC, a complex comprising Spc97, Spc98, and two Tub4 proteins ^46^. Together, Spc72 and Stu2 anchor astral microtubules at the cytoplasmic face of the SPB, resulting in accurate spindle positioning and chromosome segregation during cell division. To assess the impact of substituting Spc72, we modeled the *S. cerevisiae* Stu2 and γ-TuSC complex and replaced *S. cerevisiae* Spc72 with *K. marxianus* Spc72 *in silico*. The predicted structure revealed that *K. marxianus* Spc72 induces structural changes in *S. cerevisiae* Stu2 when these proteins are in complex (Fig. 4e). AlphaFold3 further predicted that substituting *K. marxianus* Stu2 for *S. cerevisiae* Stu2 restored the complex to a configuration resembling the wild-type *S. cerevisiae* complex (Supplementary Fig. 12). Experimental testing confirmed this prediction, showing that supplementing the strain with *K. marxianus STU2* partially rescued the incompatibility of *K. marxianus SPC72* (difference between supplemented strain vs. *S. cerevisiae* control = 79.18 ± 10.55 minutes, t-test *p* = 3.03 x 10^-15^; difference between supplemented strain vs. strain with K. marxianus *SPC72* growth defect = 59.83 ± 16.44 minutes, t-test *p* = 8.73 x 10^-11^; Fig. 4f). This partial rescue implies that incompatibility between *K. marxianus SPC72* and the *S. cerevisiae* cellular environment is more complicated than what we found for *GEM1*, likely reflecting Spc72’s dynamic interactions with other SPB proteins ^46^.

### Replacing incompatible genes completely eliminates the growth defect

Building on our findings that the coding regions of *K. marxianus GEM1* and *SPC72* drive the growth defect associated with substituting KmSeg1 for its native counterpart, we aimed to create a version of the segment that retains the maximum amount of *K. marxianus* DNA while eliminating the defect associated with substitution. To this end, we generated three constructs: one in which the coding region of *K. marxianus GEM1* was replaced with *S. cerevisiae GEM1*, another where the coding region of *K. marxianus SPC72* was replaced with *S. cerevisiae SPC72*, and a third where the coding regions of both *K. marxianus GEM1* and *SPC72* were replaced with their *S. cerevisiae* counterparts (Fig. 5a, Supplementary Fig. 7 and 13). Individually replacing *K. marxianus GEM1* or *SPC72* partially rescued the growth defect associated with substituting the *K. marxianus* segment for its native counterpart, while replacing both coding regions fully rescued it (one-way ANOVA followed by Tukey’s Honest Significant Difference tests with 95% family-wise confidence level; Fig. 5b).

**Figure 5.**
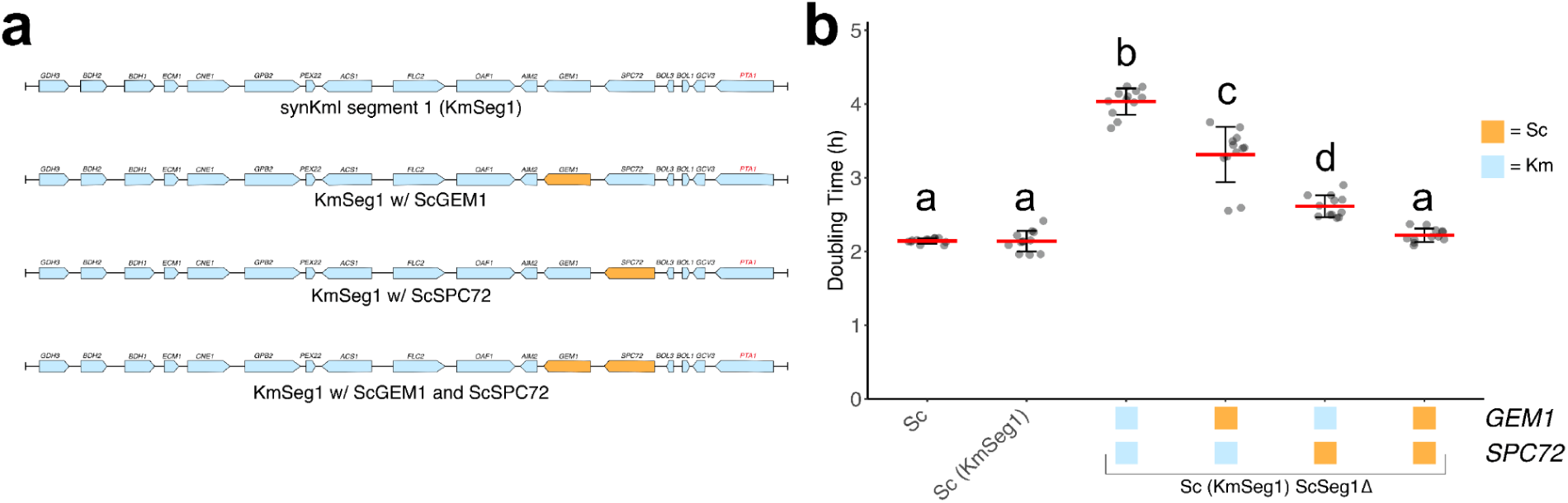
Replacing the incompatible *K. marxianus* genes on KmSeg1 completely eliminates the growth defect associated with substitution. **a.** *K. marxianus GEM1* and *SPC72* were replaced by the *S. cerevisiae* allele on the collinearized chromosome segment to produce three new variants of KmSeg1: KmSeg1 with the coding region of *K. marxianus GEM1* replaced by *S. cerevisiae GEM1*, KmSeg1 with the coding region of *K. marxianus SPC72* replaced by S. cerevisiae *SPC72*, and KmSeg1 with the coding regions of both *K. marxianus GEM1* and *SPC72* replaced with their *S. cerevisiae* counterparts. **b.** Substitution of KmSeg1 for its native counterpart resulted in a marked increase in doubling time compared to the *S. cerevisiae* control strain, indicating a significant growth defect. Individually replacing either *K. marxianus GEM1* or *K. marxianus SPC72* coding regions with their *S. cerevisiae* counterparts partially rescued the defect, resulting in doubling times that were intermediate between the KmSeg1 substitution and the *S. cerevisiae* control. Replacement of both *K. marxianus GEM1* and *SPC72* coding regions with *S. cerevisiae* alleles fully rescued the growth defect from the substitution, producing a strain with a doubling time that was indistinguishable from the *S. cerevisiae* progenitor strain (mean difference = 4.69 ± 5.87 min, one-way ANOVA followed by Tukey’s Honest Significant Difference test with 95% family-wise confidence level *p* > 0.05). Error bars represent the standard deviation from the mean. Columns sharing the same letters are not significantly different (ɑ = 0.05, One-way ANOVA and Tukey’s HSD).

In these experiments substituting KmSeg1 for its native counterpart, we observed growth effects that were similar in relative terms but less severe in absolute terms compared to the single native gene deletion experiments (Fig. 3f and 5b). These discrepancies likely reflect batch effects, as well as differences in genetic background on the effect of KmSeg1: both the entire KmSeg1 and nearly the entire native segment are present in the single gene deletions (Fig. 3f), whereas the native segment is absent in the segment substitutions (Fig. 5b). Nonetheless, the data do not provide evidence for other genes within KmSeg1 contributing to incompatibility. First, all other single native gene deletions in Fig. 3f showed the same growth rate. Second, replacing both *S. cerevisiae GEM1* and *SPC72* with their *K. marxianus* alleles produced a version of KmSeg1 that did not cause a growth defect when the native segment was deleted (Fig. 5b). We also note that most of the discrepancies we observed occurred in slow-growing strains, which may be more prone to measurement noise.

These experiments provide direct evidence for a synthetic collinearization strategy that enables the creation of a maximally *K. marxianus*-derived chromosome segment that can replace its native *S. cerevisiae* counterpart without causing growth defects.

## Discussion

Synthetic genomics offers opportunities to address longstanding biological questions that are challenging to resolve using natural systems alone. In this study, we introduced a novel application of synthetic genomics: assembling collinearized genome segments from one organism within the cellular environment of a distantly related host. Although we constructed only one-sixth of a collinear chromosome to assess feasibility, our work identifies both challenges and solutions for scaling this approach to entire chromosomes and beyond.

Collinearization overcomes key limitations that have historically restricted the genetic analysis of incompatibilities between divergent organisms, such as the inability to hybridize and produce meiotic offspring. Beyond enabling the study of reproductive isolation in existing organisms, collinearization also allows for the exploration of genetic incompatibilities between extant species and organisms that are extinct or otherwise inaccessible, as long as their genome sequence is known. In principle, the iterative replacement of all native chromosomes in a host cell with collinear chromosomes from another species could even be used in de-extinction ^47^.

Our findings support the notion that genetic incompatibilities abound between highly divergent organisms ^5–8^. Both *S. cerevisiae* and *K. marxianus* have at least 5,000 protein-coding genes. From our analysis of 17 genes, we identified two incompatible genes (∼12%). Assuming this set is representative of the entire genome, it would suggest there are at least 588 incompatibilities between these species. Among the incompatibilities we identified, both involved genes that have been annotated as nonessential, though one, *SPC72*, could be more accurately described as quasi-essential. Our findings complement a recent study that identified several essential genes causing incompatibilities between divergent yeast species ^25^. Together, these studies demonstrate that both essential and nonessential genes can play significant roles in reproductive isolation.

Our findings support the notion that incompatibilities between divergent organisms mainly stem from molecular evolution in proteins that physically interact within complexes ^25^. Coevolution within these protein complexes can lead to structural mismatches when components from divergent lineages are combined, resulting in negative epistasis. Our results indicate that these incompatibilities may vary in genetic complexity. For *GEM1*, the incompatibility appears to involve a single interacting partner, whereas for *SPC72*, it likely involves two or more. Future work should examine how a protein’s number of interaction partners affects its likelihood of being involved in incompatibilities, given the significant variability in the extent to which proteins interact ^48^.

Importantly, we also show that leading-edge software for predicting the structures of proteins within complexes has the potential to identify specific protein-protein interactions that will be incompatible between diverged organisms. Our data indicates that alleles involved in these incompatibilities will often act in a recessive manner. This implies that once incompatible proteins have been identified, supplementation with their interaction partners, as was done here and elsewhere, can mask the incompatibilities ^25^. We did not resolve the cause of the observed incompatibilities to specific amino acids, but with such resolution, it should be possible to produce substitute compatible alleles into these specific sites.

Our findings also suggest that changes in gene regulation are not a major contributor to incompatibilities between species, at least in yeasts. Despite the very high nucleotide divergence between *K. marxianus* and *S. cerevisiae* promoters, we found that most *K. marxianus* genes in KmSeg1 were transcribed in *S. cerevisiae*, though generally at different levels than their native counterparts. Moreover, substituting KmSeg1 for its native counterpart in *S. cerevisiae* caused significant changes in gene expression globally. Despite these expression differences, both within KmSeg1 and across the genome, we conclusively demonstrated that the growth defect arises directly from the coding regions of only two genes.

Understanding how genotype gives rise to phenotype is a fundamental question in the life sciences, with wide-ranging implications for evolution, inheritance, and practical applications in agriculture, biotechnology, and medicine. Collinearization provides a powerful approach for dissecting genotype-phenotype relationships, enabling research and applications in contexts where natural systems are inadequate. While our work has advanced understanding of speciation and reproductive isolation in divergent organisms, collinearization also holds significant potential for investigating other questions, such as the mechanisms behind the emergence of novel phenotypes and the genetic basis of traits with agricultural or bioengineering importance in species that cannot interbreed.

## Acknowledgements

We thank Jun-Yi Leu for input during the completion of this work and Steven Finkel for allowing us to use lab equipment. We also thank Brandon Bernardo, Matthew Dean, Suzanne Edmands, Steven Finkel, Gregory Lang, Jun-Yi Leu, Jinye Liang, Karin Pfennig, and Zachary Krieger for comments on a draft of this paper. This work was supported by NIH grants R35GM130381 and R56AI171091, NSF Designer Cells grant 2124400, a W. M. Keck Foundation Science and Engineering grant, and research funds from the University of Southern California to I.M.E, USC Student Opportunities for Academic Research and Provost’s Research Fellowships to S.H.Y., and an Agilent Postdoctoral Fellowship to A.L.V.C.

## Author Contributions

S.H.Y., A.L.V.C., and I.M.E. conceptualized this project. S.H.Y. performed the experiments. A.L.V.C., C.B.H., and D.T.L. provided technical assistance. S.H.Y. analyzed the data and produced figures. S.H.Y. and I.M.E. wrote and edited the manuscript.

## Data Availability

All data generated or analyzed in this study are included with this manuscript via the figshare link https://doi.org/10.6084/m9.figshare.29508281. Raw sequencing data for Oxford Nanopore Long-read sequencing is available in the NCBI Sequence Read archive under accession number PRJNA1284441. Raw RNA-seq data is available in the NCBI Sequence Read archive under accession number PRJNA1284543. All predicted structures for protein complexes investigated in this study are included in the figshare link as PyMol sessions (.pse) and Crystallographic Information files (.cif). Source data is also provided with this paper.

## Code Availability

Code for running software and plotting data is available via the figshare link https://doi.org/10.6084/m9.figshare.29508281.

## Methods

### Strain and genomes used for collinearization

All work done in this study was performed in the *S. cerevisiae* strain BY4742, containing a copy of the pML104 plasmid in which *URA3* had been replaced with *HIS3* ^51^. The *S. cerevisiae* genome used was S288C R64-5-1, available from the Saccharomyces Genome Database ^39^. BY4742 is a derivative of S288C. The *K. marxianus* genome used in this study was from *Kluyveromyces marxianus* DMKU3-1042, which is available under the GenBank accession identifier GCA_001417885.1 ^52^.

### Nucleotide and protein sequence divergence between the species

Orthologs between *S. cerevisiae* and *K. marxianus* were identified using only coding sequences from each species and Orthofinder2 ^53^. From the identified orthologs between *S. cerevisiae* and *K. marxianus*, only single-copy gene pairs (n = 3,788) with one-to-one ortholog relationships were selected for further analysis. Alignments for promoter regions were generated by extracting 200 bp upstream of the start codon of each gene. All alignments were performed using the Needleman-Wunsch global alignment algorithm with default parameters (gapopen = 10, gapextend = 0.5) on the EMBL-EBI Job Dispatcher ^54^. Nucleotide or protein sequence divergence was calculated as 1 - Identity(%).

### Design of synKmI

We designed synKmI in Benchling using the strategies described in the Results section and Supplementary Information. To simplify synthesis, we divided synKmI into six segments, each of which contained 200 bp of homology to its adjacent segments. The complete synKmI sequence, including its annotation, is provided as a Genbank format file with this manuscript. As discussed in the Results, we focused our synthesis and biological analyses on KmSeg1.

### Construction of KmSeg1

Twenty-five DNA fragments used for assembly were ordered from Twist Bioscience as gene fragments of ≤1.8 kb, each designed to have ∼200 bp of overlapping homology with adjacent molecules. The recipient plasmid for assembly was the BAC/YAC cloning vector pASC1, containing a *LYS2* marker for yeast and a chloramphenicol marker for *Escherichia coli* ^37^. The pASC1 vector was amplified as two PCR amplicons, each carrying one half of a split *LYS2* marker, with ∼200 bp of homology to each other and to the synthetic DNA fragments. To assemble synKmI, ∼100 ng of each DNA fragment and pASC1 piece were combined into a pool, which was then transformed into *S. cerevisiae* BY4742 using lithium acetate transformation ^55^. Transformants were plated on synthetic complete media lacking lysine and histidine (SC -His -Lys) to select for assembly. Colonies were screened for correct assembly by junction PCR across assembly junctions, followed by Oxford Nanopore Technologies sequencing.

### Oxford Nanopore Technologies sequencing of synKmI

We used long-read sequencing to identify a KmSeg1 assembly with the correct sequence and structure. Assemblies were extracted from yeast colonies using the Zymoprep Yeast Miniprep I kit (Zymo Research D2001) and electroporated into TransforMax EPI300 Electrocompetent *E. coli* (Lucigen EC300150). Transformants were recovered in SOC outgrowth medium for 1 hr with shaking and selected on LB agar plates containing chloramphenicol (30 μg/ml).

Eight bacterial colonies carrying assemblies were then inoculated into 5 mL of LB liquid media with chloramphenicol and incubated overnight at 37 °C with shaking. To increase plasmid yield, these 5 mL overnight cultures were transferred into 45 mL of LB liquid media with chloramphenicol and 50 μL of CopyControl Induction solution (LGC Biosearch Technologies), which increases the BAC copy number from 1 to 3-4 copies per cell. Cells were grown for 5 hr with shaking, and plasmid DNA was extracted using ZymoPURE II Bacterial Midiprep kit (Zymo Research). To prepare KmSeg1 assemblies for sequencing, plasmids were linearized using a single-cutter restriction enzyme AvrII.

For Oxford Nanopore Technologies library preparation, we used the manufacturer’s SQK-LSK109 and EXP-NBD104 protocols. Samples were multiplexed and sequencing was performed on a Flongle flow cell (R10.4.1) with V14 chemistry. The raw sequencing data in .pod5 format were basecalled using Oxford Nanopore Technologies’ Dorado basecaller (v4.3.0, dna_r10.4.1_e8.2_400bps_sup). Resulting fastq files were demultiplexed and barcode-trimmed in Dorado using standard parameters. Basecalled and demultiplexed fastq files were aligned to the *in silico* design of KmSeg1 using Minimap2 with default parameters ^56^. Resulting sam files were subsequently converted to BAM format, sorted, and further processed using SAMtools ^57^.

To identify sequence variants and verify sequence accuracy, we used bcftools for BCF (binary variant call format) generation ^58^. Specifically, we ran bcftools ‘mpileup’ and the resulting BCF files were processed with bcftools ‘call’ to identify any SNPs and indels relative to the reference sequence. Variant information was output as a VCF file for final analysis and a Q score threshold of 14 was applied.

### RNA-seq

Strains were grown to stationary phase overnight in SC -His -Lys at 30 °C with shaking. The next day, we used these overnights to seed new cultures in SC -His -Lys at a starting OD_600_ of ∼0.1-0.2. These setback cultures were incubated at 30 °C with shaking and harvested at mid-log phase when OD_600_ indicated that two doublings had occurred. Total RNA was extracted from these cells using the ZymoResearch YeaSTAR RNA Kit following the manufacturer’s protocol. RNA sample quality control was assessed using the Agilent 2100 Bioanalyzer System, and only RNA samples with an RNA integrity number of ≥7.5 were included. Strand-specific RNA-seq library preparation was performed by Novogene, using standard Illumina protocols. Then the libraries were sequenced by Novogene on a NovaSeqX Plus in paired-end 150 bp mode.

RNA-seq reads were aligned to the genome using Bowtie2 (v2.5.4) ^59^. Transcript quantification was performed using Kallisto and calculated as transcripts per million (TPM) ^60^. For differential analysis of gene expression data, we used sleuth ^61^. Genes were classified as upregulated or downregulated if they exhibited a log2(Fold-change) ≥ 1 and a p-value ≤ 0.05 (Wald test) with a Benjamini-Hochberg FDR threshold of 0.05. Genes that were identified to be upregulated or downregulated were input into the TheCellMap (thecellmap.org) for spatial analysis of functional enrichment ^38^. For xy scatter plots, a minimum expression threshold of transcripts per million (TPM) ≥ 1 was applied. For allele-specific expression bar plots, a pseudocount of 0.1 TPM was added before log transformation (i.e., log_2_(x + 0.1)).

### Deletion of the native S. cerevisiae chromosomal segment and genes

To perform deletions in *S. cerevisiae* BY4742, we used CRISPR-Cas9 and replaced targeted regions with a *URA3* marker. Cas9 protein was natively expressed in the cell from the pML104 plasmid containing *HIS3*. The gRNAs used for these deletions were designed *in silico* using the Benchling gRNA design tool (http://benchling.com). Only gRNAs with the highest predicted on-target scores were selected and produced by in vitro transcription, as previously described ^37^. For each gene deletion, two distinct gRNAs with high predicted on-target scores were used to ensure efficient double strand breaks. For deletion of the native counterpart of synKmI, we utilized four gRNAs in total, two targeting *GDH3* and two targeting *PTA1*. The repair templates were generated by PCR amplification of the *URA3* cassette, using tailed primers to add 40 bp of homology. For the full segment deletion, an additional 40 bp of homology was required on each side (80 bp total), which was incorporated through a second PCR using primers that annealed to the tails added in the initial cycle. Electroporation was used for these transformations, as described in ^62^. Per transformation, we used ∼600 ng of the *URA3* repair cassette and 1,200 ng of gRNAs. Transformants were selected on synthetic complete media lacking lysine, histidine, and uracil (SC -His -Lys -Ura).

### Growth assays

All phenotyping was performed on a BioTek Epoch 2 microplate spectrophotometer in 96-well plates. Yeast cells were inoculated and grown overnight under appropriate selection conditions at 30 °C with selection. 1.2 μL of the saturated yeast cultures were then inoculated into 118.8 μL of synthetic complete medium lacking lysine and histidine (SC -Lys -His). A randomized plate positioning scheme was employed to minimize potential bias across different areas of the plate. Plates were incubated at 30 °C using the double orbital shaking setting, and OD_600_ values were measured every 10 minutes until stationary phase. Doubling time values for each culture were calculated by fitting the growth data to the standard form of the logistic equation using Growthcurver ^63^. As a control, we transformed an empty pASC1 plasmid containing *LYS2* into the parental BY4742 strain with pML104 containing *HIS3*. The control strain was included in multiple replicates in every 96-well plate. For each strain, we performed at least four biological replicates, each with three technical replicates (n = 12 total measurements). All statistical tests for growth rate differences were performed in R.

### Assembly of GEM1 and SPC72 constructs

We shuffled promoter and coding sequences for *GEM1* and *SPC72* between *S. cerevisiae* and *K. marxianus*. All constructs were assembled into pASC1 containing a *LYS2* auxotrophic marker. Promoters were defined as 500 bp upstream of the start codon or the full intergenic region, whichever was shorter ^33^. For experimental consistency, *K. marxianus* terminator sequences were used across all chimeric constructs, here defined as 100 bp downstream of the stop codon. Promoters and coding regions were ordered separately as synthetic DNA fragments from Twist, and assembled with each other and pASC1 in yeast by electroporation ^62^. Fragments contained ≥200 bp overlapping homology to pASC1 at one end and homology to a promoter or coding region at the other end. As we did for KmSeg1, the pASC1 vector was amplified as two PCR amplicons with split *LYS2* markers, each with ∼200 bp of overlapping homology to one another and to synthetic DNA fragments. Transformants were selected on synthetic media lacking lysine and histidine (SC -His -Lys). All *GEM1* and *SPC72* constructs were verified by Oxford Nanopore Technologies sequencing with Plasmidsaurus.

### Supplementation experiments

Supplementation plasmids were made that each contained two interacting genes: *GEM1* and *MMM1* (*GEM1-MMM1*) or *SPC72* and *STU2* (*SPC72-STU2*). These constructs were generated by ordering fragments containing the *S. cerevisiae* promoter for these genes and the *K. marxianus* coding regions. These fragments were obtained from Twist Bioscience and assembled into pASC1 containing *LYS2* using yeast electroporation ^62^. All constructs were verified by Oxford Nanopore Technologies sequencing with Plasmidsaurus. The *GEM1-MMM1* construct was transformed into *S. cerevisiae* BY4742 *gem1*Δ. The *SPC72-STU2* construct was transformed into *S. cerevisiae* BY4742, and the endogenous copy of *SPC72* was deleted afterward, as we were unable to recover an *SPC72* deletion in wild type *S. cerevisiae*.

### Protein complex interactions

All protein complexes investigated in this study were predicted using AlphaFold3 through the AlphaFold server (https://alphafoldserver.com/) ^42^. We used the PyMol visualization software (Schrodinger) to visualize predicted protein complex structures ^64^.

### A fully compatible KmSeg1

To produce a version of KmSeg1 that is fully compatible with an *S. cerevisiae* cellular environment, we replaced the *K. marxianus GEM1* and *SPC72* coding regions with alleles from *S. cerevisiae*. To make these changes, we used an editing strategy performed in yeast. For each gene, we designed two gRNAs to target the *K. marxianus* coding region, along with a repair cassette containing the *S. cerevisiae* coding region flanked by homology to the *K. marxianus* segment. A pRS316 plasmid, which contains *URA3*, was cotransformed along with the repair cassette and gRNAs using standard yeast electroporation to select for transformants on synthetic complete medium lacking lysine, histidine, and uracil (SC -His -Lys -Ura). Colonies were screened by PCR for correct replacement of *GEM1* or *SPC72* in KmSeg1 and then further verified Sanger sequencing through the entire replaced gene (GeneWiz). To induce loss of the pRS316, colonies with the correct constructs were plated on media containing 5-Fluoroorotic acid (5-FOA) ^65^.

## References

1. Carroll, S. B., Greiner, J. K. & Weatherbee, S. D. From DNA to Diversity: Molecular Genetics and the Evolution of Animal Design. (Blackwell, Malden, MA, 2004).

2. Presgraves, D. C. The molecular evolutionary basis of species formation. Nat. Rev. Genet. 11, 175–180 (2010).

3. Lin, R.-C., Ferreira, B. T. & Yuan, Y.-W. The molecular basis of phenotypic evolution: beyond the usual suspects. Trends Genet. 40, 668–680 (2024).

4. Lynch, M. & Walsh, B. Genetics and Analysis of Quantitative Traits. (Oxford University Press, Oxford, New York, 1998).

5. Orr, H. A. The population genetics of speciation: the evolution of hybrid incompatibilities. Genetics 139, 1805–1813 (1995).

6. Orr, H. A. & Turelli, M. The evolution of postzygotic isolation: accumulating Dobzhansky-Muller incompatibilities. Evol. Int. J. Org. Evol. 55, 1085–1094 (2001).

7. Matute, D. R., Butler, I. A., Turissini, D. A. & Coyne, J. A. A Test of the Snowball Theory for the Rate of Evolution of Hybrid Incompatibilities. Science 329, 1518–1521 (2010).

8. Maheshwari, S. & Barbash, D. A. The Genetics of Hybrid Incompatibilities. Annu. Rev. Genet. 45, 331–355 (2011).

9. Coradini, A. L. V., Hull, C. B. & Ehrenreich, I. M. Building genomes to understand biology. Nat. Commun. 11, 6177 (2020).

10. James, J. S., Dai, J., Chew, W. L. & Cai, Y. The design and engineering of synthetic genomes. Nat. Rev. Genet. 1–22 (2024) doi:10.1038/s41576-024-00786-y.

11. Hutchison, C. A. et al. Design and synthesis of a minimal bacterial genome. Science 351, aad6253 (2016).

12. Fredens, J. et al. Total synthesis of Escherichia coli with a recoded genome. Nature 569, 514–518 (2019).

13. Pinglay, S. et al. Synthetic regulatory reconstitution reveals principles of mammalian Hox cluster regulation. Science 377, eabk2820 (2022).

14. Delneri, D. et al. Engineering evolution to study speciation in yeasts. Nature 422, 68–72 (2003).

15. Richardson, S. M. et al. Design of a synthetic yeast genome. Science 355, 1040–1044 (2017).

16. Hunter, N., Chambers, S. R., Louis, E. J. & Borts, R. H. The mismatch repair system contributes to meiotic sterility in an interspecific yeast hybrid. EMBO J. 15, 1726–1733 (1996).

17. Greig, D., Louis, E. J., Borts, R. H. & Travisano, M. Hybrid Speciation in Experimental Populations of Yeast. Science 298, 1773–1775 (2002).

18. Liti, G., Barton, D. B. H. & Louis, E. J. Sequence Diversity, Reproductive Isolation and Species Concepts in Saccharomyces. Genetics 174, 839–850 (2006).

19. Lee, H.-Y. et al. Incompatibility of Nuclear and Mitochondrial Genomes Causes Hybrid Sterility between Two Yeast Species. Cell 135, 1065–1073 (2008).

20. Hou, J., Friedrich, A., de Montigny, J. & Schacherer, J. Chromosomal Rearrangements as a Major Mechanism in the Onset of Reproductive Isolation in *Saccharomyces cerevisiae*. Curr. Biol. 24, 1153–1159 (2014).

21. Hou, J., Friedrich, A., Gounot, J.-S. & Schacherer, J. Comprehensive survey of condition-specific reproductive isolation reveals genetic incompatibility in yeast. Nat. Commun. 6, 7214 (2015).

22. Leducq, J.-B. et al. Speciation driven by hybridization and chromosomal plasticity in a wild yeast. Nat. Microbiol. 1, 1–10 (2016).

23. Jhuang, H., Lee, H. & Leu, J. Mitochondrial–nuclear co-evolution leads to hybrid incompatibility through pentatricopeptide repeat proteins. EMBO Rep. 18, 87–101 (2017).

24. Swamy, K. B. S. et al. Proteotoxicity caused by perturbed protein complexes underlies hybrid incompatibility in yeast. Nat. Commun. 13, 4394 (2022).

25. Lai, H.-Y., Yu, Y.-H., Jhou, Y.-T., Liao, C.-W. & Leu, J.-Y. Multiple intermolecular interactions facilitate rapid evolution of essential genes. Nat. Ecol. Evol. 7, 745–755 (2023).

26. Karim, A., Gerliani, N. & Aïder, M. Kluyveromyces marxianus: An emerging yeast cell factory for applications in food and biotechnology. Int. J. Food Microbiol. 333, 108818 (2020).

27. Wolfe, K. H. & Shields, D. C. Molecular evidence for an ancient duplication of the entire yeast genome. Nature 387, 708–713 (1997).

28. Shen, X.-X. et al. Tempo and Mode of Genome Evolution in the Budding Yeast Subphylum. Cell 175, 1533–1545.e20 (2018).

29. Kellis, M., Birren, B. W. & Lander, E. S. Proof and evolutionary analysis of ancient genome duplication in the yeast Saccharomyces cerevisiae. Nature 428, 617–624 (2004).

30. Yue, J.-X. et al. Contrasting evolutionary genome dynamics between domesticated and wild yeasts. Nat. Genet. 49, 913–924 (2017).

31. Albonico, F., B., E., G, P. H. & B., D. New Saccharomyces cerevisiae-Kluyveromyces marxianus fusant shows enhanced alcoholic fermentation performance. World J. Microbiol. Biotechnol. 38, 251 (2022).

32. Lyu, Y. et al. Transferring functional chromosomes between intergeneric yeast generates monochromosomal hybrids with improved phenotypes and widespread transcriptional responses. 2025.01.24.634819 Preprint at 10.1101/2025.01.24.634819 (2025).

33. Guo, Y. et al. YeastFab: the design and construction of standard biological parts for metabolic engineering in Saccharomyces cerevisiae. Nucleic Acids Res. 43, e88 (2015).

34. Graber, J. H., McAllister, G. D. & Smith, T. F. Probabilistic prediction of Saccharomyces cerevisiae mRNA 3′-processing sites. Nucleic Acids Res. 30, 1851–1858 (2002).

35. Curran, K. A. et al. Short Synthetic Terminators for Improved Heterologous Gene Expression in Yeast. ACS Synth. Biol. 4, 824–832 (2015).

36. Altschul, S. F., Gish, W., Miller, W., Myers, E. W. & Lipman, D. J. Basic local alignment search tool. J. Mol. Biol. 215, 403–410 (1990).

37. Coradini, A. L. V. et al. Building synthetic chromosomes from natural DNA. Nat. Commun. 14, 8337 (2023).

38. Usaj, M. et al. TheCellMap.org: A Web-Accessible Database for Visualizing and Mining the Global Yeast Genetic Interaction Network. G3 GenesGenomesGenetics 7, 1539–1549 (2017).

39. Engel, S. R. et al. Saccharomyces Genome Database: advances in genome annotation, expanded biochemical pathways, and other key enhancements. Genetics iyae185 (2024) doi:10.1093/genetics/iyae185.

40. Kornmann, B., Osman, C. & Walter, P. The conserved GTPase Gem1 regulates endoplasmic reticulum–mitochondria connections. Proc. Natl. Acad. Sci. 108, 14151–14156 (2011).

41. Souès, S. & Adams, I. R. SPC72: a spindle pole component required for spindle orientation in the yeast Saccharomyces cerevisiae. J. Cell Sci. 111, 2809–2818 (1998).

42. Abramson, J. et al. Accurate structure prediction of biomolecular interactions with AlphaFold 3. Nature 630, 493–500 (2024).

43. Michel, A. H. & Kornmann, B. The ERMES complex and ER–mitochondria connections. Biochem. Soc. Trans. 40, 445–450 (2012).

44. Dobzhansky, T. Studies on hybrid sterility. Z. Für Zellforsch. Mikrosk. Anat. 21, 169–223 (1934).

45. Muller, H. J. Isolating mechanisms, evolution, and temperature. Biol. Symp. 6, 71–125 (1942).

46. Usui, T., Maekawa, H., Pereira, G. & Schiebel, E. The XMAP215 homologue Stu2 at yeast spindle pole bodies regulates microtubule dynamics and anchorage. EMBO J. 22, 4779–4793 (2003).

47. Sherkow, J. S. & Greely, H. T. What If Extinction Is Not Forever? Science 340, 32–33 (2013).

48. Michaelis, A. C. et al. The social and structural architecture of the yeast protein interactome. Nature 624, 192–200 (2023).

49. Kurtzman, C. P. & Robnett, C. J. Phylogenetic relationships among yeasts of the ‘Saccharomyces complex’ determined from multigene sequence analyses. FEMS Yeast Res. 3, 417–432 (2003).

50. Hagman, A., Säll, T., Compagno, C. & Piskur, J. Yeast “Make-Accumulate-Consume” Life Strategy Evolved as a Multi-Step Process That Predates the Whole Genome Duplication. PLOS ONE 8, e68734 (2013).

51. Laughery, M. F. et al. New vectors for simple and streamlined CRISPR-Cas9 genome editing in *Saccharomyces cerevisiae*: Vectors for simple CRISPR-Cas9 genome editing in yeast. Yeast 32, 711–720 (2015).

52. Lertwattanasakul, N. et al. Genetic basis of the highly efficient yeast Kluyveromyces marxianus: complete genome sequence and transcriptome analyses. Biotechnol. Biofuels 8, 47 (2015).

53. Emms, D. M. & Kelly, S. OrthoFinder: phylogenetic orthology inference for comparative genomics. Genome Biol. 20, 238 (2019).

54. Madeira, F. et al. The EMBL-EBI Job Dispatcher sequence analysis tools framework in 2024. Nucleic Acids Res. 52, W521–W525 (2024).

55. Gietz, R. D. & Schiestl, R. H. High-efficiency yeast transformation using the LiAc/SS carrier DNA/PEG method. Nat. Protoc. 2, 31–34 (2007).

56. Li, H. Minimap2: pairwise alignment for nucleotide sequences. Bioinformatics 34, 3094–3100 (2018).

57. Li, H. et al. The Sequence Alignment/Map format and SAMtools. Bioinformatics 25, 2078–2079 (2009).

58. Li, H. A statistical framework for SNP calling, mutation discovery, association mapping and population genetical parameter estimation from sequencing data. Bioinformatics 27, 2987–2993 (2011).

59. Langmead, B. & Salzberg, S. L. Fast gapped-read alignment with Bowtie 2. Nat. Methods 9, 357–359 (2012).

60. Bray, N. L., Pimentel, H., Melsted, P. & Pachter, L. Near-optimal probabilistic RNA-seq quantification. Nat. Biotechnol. 34, 525–527 (2016).

61. Pimentel, H., Bray, N. L., Puente, S., Melsted, P. & Pachter, L. Differential analysis of RNA-seq incorporating quantification uncertainty. Nat. Methods 14, 687–690 (2017).

62. Kannan, K. et al. One step engineering of the small-subunit ribosomal RNA using CRISPR/Cas9. Sci. Rep. 6, 30714 (2016).

63. Sprouffske, K. & Wagner, A. Growthcurver: an R package for obtaining interpretable metrics from microbial growth curves. BMC Bioinformatics 17, 172 (2016).

64. The PyMOL Molecular Graphics System. Schrödinger, LLC.

65. Boeke, J. D., Trueheart, J., Natsoulis, G. & Fink, G. R. [10] 5-Fluoroorotic acid as a selective agent in yeast molecular genetics. in Methods in Enzymology vol. 154 164–175 (Academic Press, 1987).

